# Persistence and *in vivo* evolution of vaginal bacterial strains over a multi-year time period

**DOI:** 10.1101/2022.04.27.489821

**Authors:** Michael France, Bing Ma, Jacques Ravel

## Abstract

It is not clear if the bacterial strains which comprise our microbiota are mostly long-term colonizers or transient residents. Studies have demonstrated decades long persistence of bacterial strains within the gut, but persistence at other body sites has yet to be determined. The vaginal microbiota (VMB) is often dominated by *Lactobacillus*, although it is also commonly comprised of a more diverse set of other facultative and obligate anaerobes. Longitudinal studies have demonstrated that these communities can be stable over several menstrual cycles or can fluctuate temporally in species composition. We sought to determine whether the bacterial strains which comprise the VMB were capable of persisting over longer time-periods. We performed shotgun metagenomics on paired samples from 10 participants collected 1 and 2 years apart. The resulting sequences were *de novo* assembled and binned into high-quality metagenome assembled genomes. Persistent strains were identified based on the sequence similarity between the genomes present at the two timepoints and were found in the VMB of six of the participants, three of which had multiple. The VMB of the remaining four participants was similar in species composition at the two timepoints but was comprised of different strains. For the persistent strains, we were able to identify the mutations which fixed in the populations over the observed time period, giving insight into the evolution of these bacteria. These results indicate that bacterial strains can persist in the vagina for extended periods of time, providing an opportunity for them to evolve in the host microenvironment.

**Importance:** The persistence of strains within the vaginal microbiota has not yet been characterized. Should these strains be capable of persisting for extended periods of time, they could evolve within their host in response to selective pressures exerted by the host or by other members of the community. Here, we present preliminary findings which demonstrate that bacterial strains can persist in the vagina for at least one year and evolve over time. In several cases, multiple strains persisted together in a community, indicating that co-evolution between bacterial strains could occur in the vagina. Our observations motivate future studies which collect samples from more participants, at more timepoints and over even longer periods of time. Understanding which strains persist, what factors drive their persistence, and what selective pressures they face will inform the development and delivery of rationally designed live biotherapeutics for the vagina.

## Observation

The human microbiome is estimated to be comprised of hundreds to thousands of distinct bacterial species and strains (1, 2). These bacteria often live in close association with host tissues and are thought to be critical for the maintenance of our health (3). Studies on the gut microbiome have demonstrated that strains of these species are capable of persisting within a host for extended periods of time (4, 5). However, the potential for long-term persistence of bacterial strains at other body sites remains largely unexplored. The microbial communities which inhabit the vagina are unique from those found at other body sites (6). They are often dominated (>90% relative abundance) by single species of *Lactobacillus*, although a significant proportion of women have more compositionally even communities containing an assortment of facultative and obligate anaerobic bacteria (7, 8). Communities which are dominated by *Lactobacillus* spp. have been associated with a decreased risk for several adverse health outcomes, leading many to consider them to be “optimal” (reviewed in M. France et al. (9)). Observational studies have demonstrated that the vaginal microbiota (VMB) of some women maintain species composition over several menstrual cycles, while others have communities which vary over time (10, 11). It has yet to be determined if the VMB typically maintains bacterial strains composition over longer-periods of time or if there are frequent turnovers in the dominant strain of each species.

In this study, we sought to characterize the long-term persistence and *in vivo* evolution of bacterial strains within the VMB. Women whose communities had similar species composition at timepoints separated by at least one year were identified using previously published 16S rRNA gene amplicon survey data (7, 11) and ten were selected to represent the breadth of commonly observed community compositions. Shotgun metagenomes were generated to characterize the strains present at each timepoint (12). The resulting sequence reads were mapped to the VIRGO non-redundant gene catalog (13) to establish the taxonomic composition of each sample (**Figure 1A**). All participants had similar species in their VMB at the two timepoints but in the cases of participants 4 & 5, their relative abundances had shifted substantially. Both of these participants had communities that were dominated by *L. iners* at the initial timepoint which was later supplanted by either *L. crispatus* (participant 4) or *L. jensenii* (participant 5).

**Figure 1:**
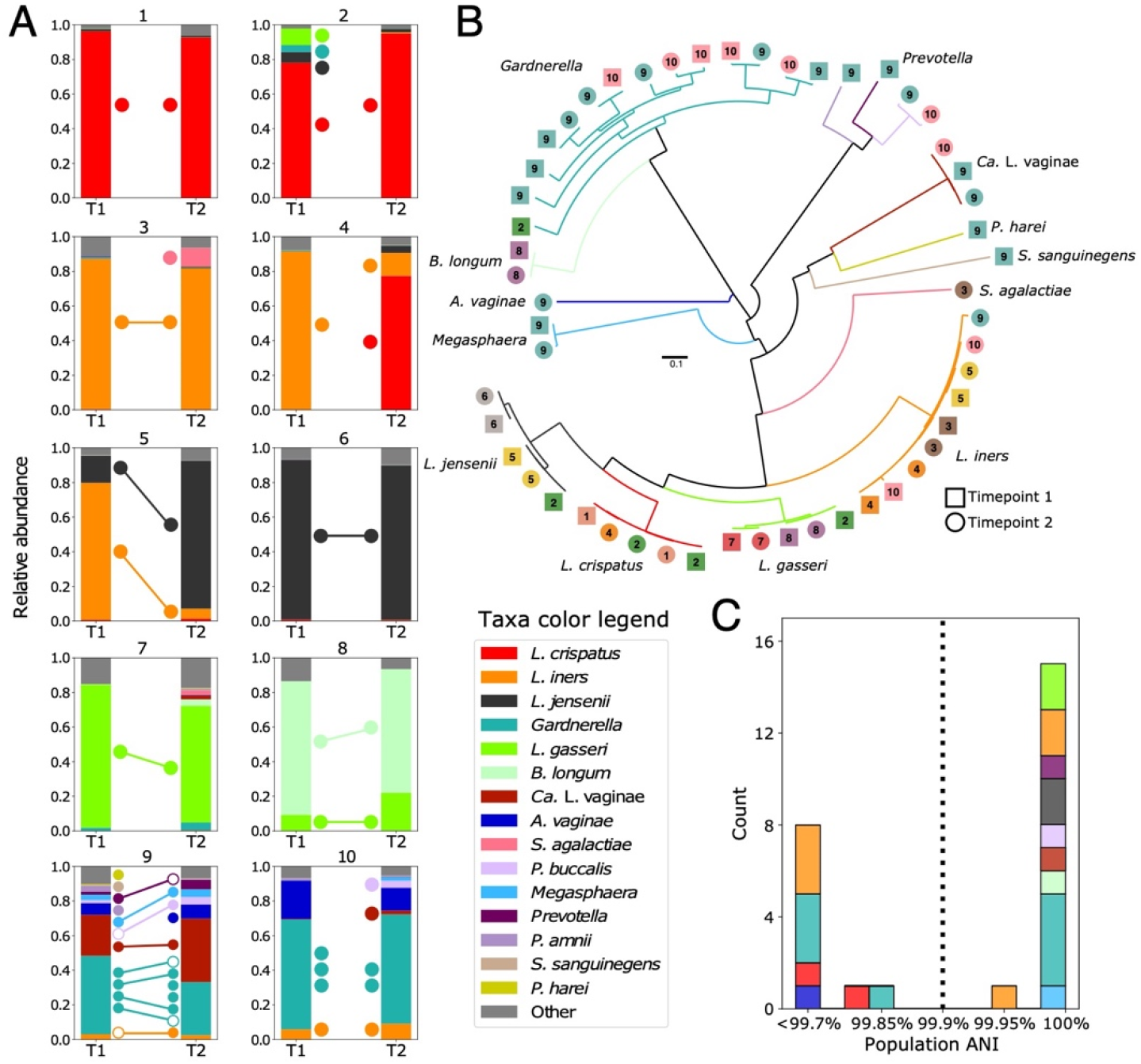
Taxonomic composition of each participant’s VMB at the two timepoints as established using the VIRGO non-redundant gene catalog. (**A**) Points between the bars represent the MAGs recovered from each timepoint (closed) or strains identified by inStrain but not assembled (open). Where the two points are connected by a line, the strain met the 99.9% ANI threshold and was considered to be present at both timepoints. Phylogenetic tree was derived from a concatenated alignment of 100 orthologous genes identified in at least 98% of the 53 bacterial metagenome assembled genomes (MAGs). (**B**) Branches are colored according to taxonomy and MAGs are labeled with the participant number and the timepoint (square-T1, circle-T2). Average nucleotide identity (ANI) between strains of the same species that were identified at both timepoints for a participant. (**C**) An ANI of at least 99.9% was considered the threshold for strain persistence in a participant’s VMB. Stacked bars are colored according to taxonomy.

We next sought to determine which participants had maintained the same strain(s) over the 1-2-year time period. *De novo* assembly using metaSPAdes (14) and contig binning, as described previously (15), produced 53 metagenome assembled genomes (MAGs), representing 15 species (**Figure 1B**). To identify which participants had the same strain(s) at the two timepoints, we used inStrain (16), with a percent identify threshold of at least 99.9%. This relatively strict threshold was chosen as any greater degree of sequence divergence would be difficult to explain given estimated substitution rates for bacteria (17). Sixteen strains representing nine species were identified at both timepoints from a single individual (**Figure 1C**). We conclude that these observations result from the persistence of the strain(s) within a participant’s VMB. Six of the ten participants were found to have at least one persistent strain in their VMB, and three (participants 5, 8 & 9) had multiple (**Figure 1A**). Of those, two (participants 5 & 8) had communities which were primarily comprised of two species whose strains had persisted (participant 5: *L. iners* and *L. jensenii*, participant 8: *B. longum* and *L. gasseri*). Participant 9 had a more diverse VMB that was not dominated by *Lactobacillus* spp. and was found to have nine persistent strains including four strains of *Gardnerella*, two strains of *Prevotella* and one strain each of: *L. iners, Ca*. L. vaginae, and *Megasphaera*. This observation indicates that strain persistence is not just a property of *Lactobacillus* dominant communities and that these more compositionally even communities can also exhibit long-term stability in strain composition.

The remaining four participants were not found to have any persistent strains, despite the similarity in their taxonomic composition at the two timepoints. This included the two participants that had a *L. crispatus* dominant VMB at both timepoints (participants 1 & 2, **Figure 1A**), indicating that the *L. crispatus* at the initial timepoint was supplanted by another at the second sampling. It could be that these *L. crispatus* populations went extinct and were subsequently reestablished, or that the population experienced a shift in the dominant strain, as prior studies have indicated these populations are often comprised of multiple strains (13, 18). Participant 4 had a *L. iners* dominated VMB at the initial timepoint, which shifted to a community which contained a majority of *L. crispatus* and a minority of *L. iners* at the second timepoint. The *L. iners* identified at the second timepoint was not the same strain as that identified at the initial timepoint. Finally, participant 10 had the more diverse VMB at both timepoints with similar species composition, but, unlike participant 9, was not found to have any persistent strains. These observations demonstrate that consistency in species composition in a VMB over time does not necessarily reflect the persistence of individual strains.

Long-term colonization of a strain in the VMB provides an opportunity for the strain’s population to adapt to a specific host environment. For the six participants with persistent strains, we were able to identify mutations that had occurred and been fixed in the population. BreSeq was used to characterize genomic changes in nine of the persistent strains with sufficient coverage (19). The substitutions were observed in a variety of genetic loci and included nonsynonymous and synonymous changes as well as small insertions or deletions (summarized in **Table 1**, details in **Supplemental table 1**). The average number of substitutions observed for each strain was ~16 providing an average substitution rate of 7.74×10^−6^ substitutions per bp per year. These observed substitution rates are on the higher end of previous estimates for bacteria which range from 10^−8^ to 10^−5^ substitutions per bp per year (17, 20). The observed high substitution rate could be explained by an increased mutation rate in vaginal bacteria as most have undergone substantial genome reduction, which is often accompanied by a loss of DNA repair elements (21). Some of these mutations, chiefly those that result in an amino acid change, could be adaptive, although it is impossible to discern without additional evidence. Observations at intervening timepoints could inform the order and speed of each fixation event, and reveal which mutations occurred on the same background and fixed jointly. Additionally, observing the long-term strain persistence and evolution in a larger cohort could reveal instances of parallel evolution, indicating an adaptive role.

**Table 1:** Mutations identified in persistent strains and corresponding estimates of the substitution rate

| Participant | Species | No. Mutations* | Genes with mutations** | Subst. Rate† |
| --- | --- | --- | --- | --- |
| 3 | *L. iners* | 13 (5/5/0/3) | **wecH**, DagR, **sbnD**, **yheH**, mgIA, **recF** | 5.4x10<sup>-6</sup> |
| 5 | *L. iners* | 15 (7/1/2/5) | **adk**, **spxA**, rfbX, **btuD**, **dhaL** | 6.5x10<sup>-6</sup> |
| 5 | *L. jensenii* | 10 (6/2/1/1) | **yheL**, **dedA**, **ebh**, **arcD1**, **mutS2** | 3.4x10<sup>-6</sup> |
| 6 | *L. jensenii* | 13 (3/2/5/3) | yajL, glf, citX, glvR | 6.4x10<sup>-6</sup> |
| 7 | *L. gasseri* | 10 (1/0/3/6) | **nrdD**, **hpt**, **cls** | 3.6x10<sup>-6</sup> |
| 8 | *B. longum* | 22 (6/4/1/11) | **degA**, gndA, **thiC**, nusB, **hadI**, **glgB**, acn | 6.4x10<sup>-6</sup> |
| 9 | *Ca. L. vaginae* | 8 (2/5/0/1) | IsaC | 5.0x10<sup>-6</sup> |
| 9 | *Gardnerella gsp 7* | 15 (0/11/0/4) | glgE, gatB, thiO, carA, ykoE, ybhL | 1.1x10<sup>-5</sup> |
| 9 | *G. vaginalis* | 39 (7/29/0/3) | malP,gla, trmH,gatA,lysS,infC,thiO,prfA,**tuf**,**glgX**,tal,tkt,msbA,uvrB,accA | 2.7x10<sup>-5</sup> |
| *Number of Mutations (nonsynonymous;synonymous;indel;intergenic/noncoding) |  |  |  |  |
| **Genes with mutations, mutations in bolded genes caused a change in amino acid sequence, hypothetical genes not shown |  |  |  |  |
| †Units: mutations per bp per year |  |  |  |  |

## Conclusion

We observed the persistence of bacterial strains in the vaginal microbiota over a one-to-two-year period. Several participants had multiple persistent bacterial strains, opening the window for coevolution to occur between cohabiting strains. It is not clear why some participants maintained their strains, while others did not. Host factors are expected to play a principal role and would include things like the use of antibiotics or the introduction of novel sexual partners (22). However, it could also be that some species or strains are more capable long-term colonizers of the vaginal niche than others and that microbial factors play a disproportionate role. Larger studies are needed to characterize the determinants of strain persistence in the vaginal microbiota. For the persistent strains we were also able to characterize the mutations which occurred and reached fixation over this time period. These observations provide an initial glimpse into the *in vivo* evolution of the vaginal microbiota and inform expected rates of change for these bacteria. From these preliminary data, we conclude that the strain composition of the vaginal microbiota is often stable over long periods of time but should not be assumed.

## Data availability

Shotgun metagenome data have been deposited in the Short Read Archive: <u>PRJNA575586</u>. All scripts used in the processing and analyses of the metagenomes are available at: https://github.com/ravel-lab/two_year.

## Acknowledgements

This work was supported in part by the National Institute of Allergy and Infectious Diseases and the National Institute for Nursing Research of the National Institutes of Health under award numbers UH2AI083264 and R01NR015495, respectively.

## Competing interests

J.R. is a cofounder of LUCA Biologics, a biotechnology company focusing on translating microbiome research into live biotherapeutic drugs for women’s health.

All other authors declare no competing interests.

## Supplemental information

**Supplemental text 1**: Experimental methods. Detailed description of the experimental and bioinformatic procedures used to generate and analyze the shotgun metagenomics data.

**Supplemental table 1**: Description of mutations which fixed in the populations of persistent strains

